# Hydrodynamics structure the physiological trajectory and bloom dynamics of a harmful diatom species across neighboring coastal ecosystems

**DOI:** 10.64898/2026.10.05.756714

**Authors:** Mickael Le Gac, Léa Prigent, Julien Quéré, Florian Caradec, Fabienne Hervé, Emilie Rabiller, Zouher Amzil, Martin Plus

## Abstract

Hydrodynamics is a strong regulator of phytoplankton cell densities and population connectivity. However, the extent to which hydrodynamics, specifically dilution and advection, dictates the internal physiological state and overall bloom dynamics remains poorly understood. We conducted an integrated study of the harmful diatom *Pseudo-nitzschia australis* in two neighboring coastal bays with similar abiotic conditions but contrasting bloom trajectories. Using a combination of field observations, transcriptomics, domoic acid (DA) measurements, and hydrodynamic simulations, we show that a three-fold difference in dilution rates alone explains a 25-day shift in bloom timing and an order-of-magnitude difference in cell densities. Transcriptomic profiling revealed distinct metabolic and stress-associated gene modules that synchronized with hydrodynamically-mediated nutrient depletion rate, linking residence times to cellular mortality. Furthermore, directional connectivity between the bays may explain population-level toxin heterogeneity, as advection of older cells, releasing DA outside the cells, may have decreased the proportion of intracellular DA in the destination bay. Our results illustrate that hydrodynamics do not merely modulate population transport; they structure the full physiological life cycle of a bloom, from initiation to nutrient-driven demise. This highlights the necessity of considering hydrodynamics as a primary explanation for bloom dynamics with cascading effects on bloom onset, cell density, nutrient consumption, cell physiology and mortality.

## Introduction

Phytoplankton blooms are fundamental drivers of global biogeochemical cycles and coastal food web dynamics (Falkowski et al.; 2008; Carstensen et al., 2015). The mechanisms that control the timing, intensity, and termination of these blooms are not fully understood (Carstensen et al. 2015; Caracciolo et al. 2022). Traditionally, bloom variability has been attributed to differences in light, temperature, nutrient availability, grazing pressure, or other local ecological controls (Carstensen et al., 2015). Hydrodynamic forcing is also a primary architect of these events. At the mesoscale, hydrodynamic features strongly influence the dispersal, patchiness, and dilution of phytoplankton biomass (Lévy et al., 2024; Lehahn et al., 2017; Ser-Giacomi et al., 2023). Vertical processes such as mixed-layer deepening and upwelling are indirect regulators by modulating light exposure and nutrient supply (Behrenfeld and Boss, 2017; Diaz et al., 2021; Pitcher et al. 2010). In coastal ecosystems, the succession of spring and neap tides often induce biomass accumulation in stratified waters during neap tides and biomass dilution during more energetic spring tides (Cloern 1991; Blauw et al., 2012; Cadier et al., 2017).

In coastal ecosystems, phytoplankton blooms often exhibit strong spatio-temporal heterogeneity, with large shifts in magnitude and timing occurring over distances of just a few kilometers (Song et al., 2019; Ferreira et al., 2021). A hydrodynamic forcing that appears spatially uniform rarely produces a uniform hydrodynamic response at the coast. For instance, tidal currents interacting with complex topography and bathymetry generate current fields that differ sharply between nearby locations, producing pronounced spatial heterogeneity in residence time and flushing rates (Signel and Geyer 1991; Monsen et al., 2002). Coastline geometry can also generate retentive recirculation zones immediately adjacent to rapidly flushed waters during wind-driven upwelling events (Largier, 2020). The biological consequences of this heterogeneity are well-documented for cell density and biomass (Wan et al., 2013; Qin and Chen, 2019; Ralston et al., 2016; Diaz et al., 2025). However, it remains unclear how these physical contrasts influence the finer-scale physiological state of the populations.

Metatranscriptomics has recently emerged as a powerful tool for decoding these physiological signals directly in situ. The analyses of environmental mRNA identified molecular evidence for resource partitioning among co-occurring diatoms (Alexander et al., 2015), specific responses to iron limitation (Moreno et al., 2024) or light-driven metabolic shifts during mixed-layer shallowing (Diaz et al., 2023). Recent studies have also begun to link molecular signatures to life-cycle events, such as synchronized sexual reproduction during a *Pseudo-nitzschia australis* bloom (Prigent et al., 2025). Collectively, these approaches demonstrate that metatranscriptomics can diagnose the precise physiological state of a microbial population.

Diatoms of the genus *Pseudo-nitzschia* are cosmopolites and may represent 5% of the diatoms worldwide (Malviya et al., 2016). More than 50 species have been described, of which about half have been documented as producing domoic acid (DA), a potent neurotoxin that bioaccumulates in food webs and may cause Amnesic Shellfish Poisoning (ASP) (Bates et al., 2018; Trainer et al., 2012). Among these, *P. australis* is particularly problematic, forming blooms that pose severe risks to human health and food security (McCabe et al., 2016; Brunson et al., 2024). In the Northeast Atlantic, *P. australis* blooms are characterized by high spatial variability, with neighboring bays exhibiting divergent bloom timing and magnitudes under similar regional forcing (Belin et al., 2023). Here, we utilize this system to test whether physical residence time and connectivity act as the primary drivers of microbial physiological trajectories. By coupling realistic hydrodynamic simulations with metatranscriptomics and toxin analysis, we demonstrate that hydrodynamics does not simply modulate population transport, but defines the entire molecular life history of the bloom from initiation to toxin phenotype and physiological decline.

## 2. Materials and Methods

### 2.1. Study area

The study site is on the north-east coast of the Atlantic ocean in North-Western France. It is a macrotidal (maximum tidal amplitude 8 m, tide periodicity: 12 h) coastline where the bays of Brest (180 km^2^) and Douarnenez (350 km^2^) open up and connect with the Iroise Sea (North-East Atlantic). The Bay of Brest is a semi-enclosed embayment connected to shelf waters by a narrow strait (2 km wide, 40 m deep; Le Pape et al., 1996). The two main freshwater inputs in the bay are the Aulne and Elorn rivers (average annual flow 21 and 5 m^3^ s^−1^, respectively). In this bay tidal currents and wind ensure a quasi-permanent vertical mixing of the waters (Beucher et al., 2004). The Bay of Douarnenez can experience short thermal stratification periods during calm summer periods and receive much lower riverine inputs with mean flows below 2 m^3^.s^-1^ (Merceron et al., 2007; Guillam et al., 2020).

Following initial detection of the toxic *P. australis* species by the French phytoplankton monitoring network (REPHY), a total of 60 water samples were collected bi-weekly from shore between February and April 2021 at four monitoring stations (Fig. 1, Supplementary material 1). Two monitoring stations are located in the Bay of Brest (Lanvéoc: 48.2929°N; -4.4584°E, Sainte-Anne: 48.3601°N; -4.5531°E), one monitoring station is located in the Bay of Douarnenez (Telgruc station: 48.2099°N, -4.3733°E) and the last one is located between the other three in Dinan-Kerloc’h and is more open to the Iroise Sea (48.2570°N; -4.5609°E)

**Figure 1.**
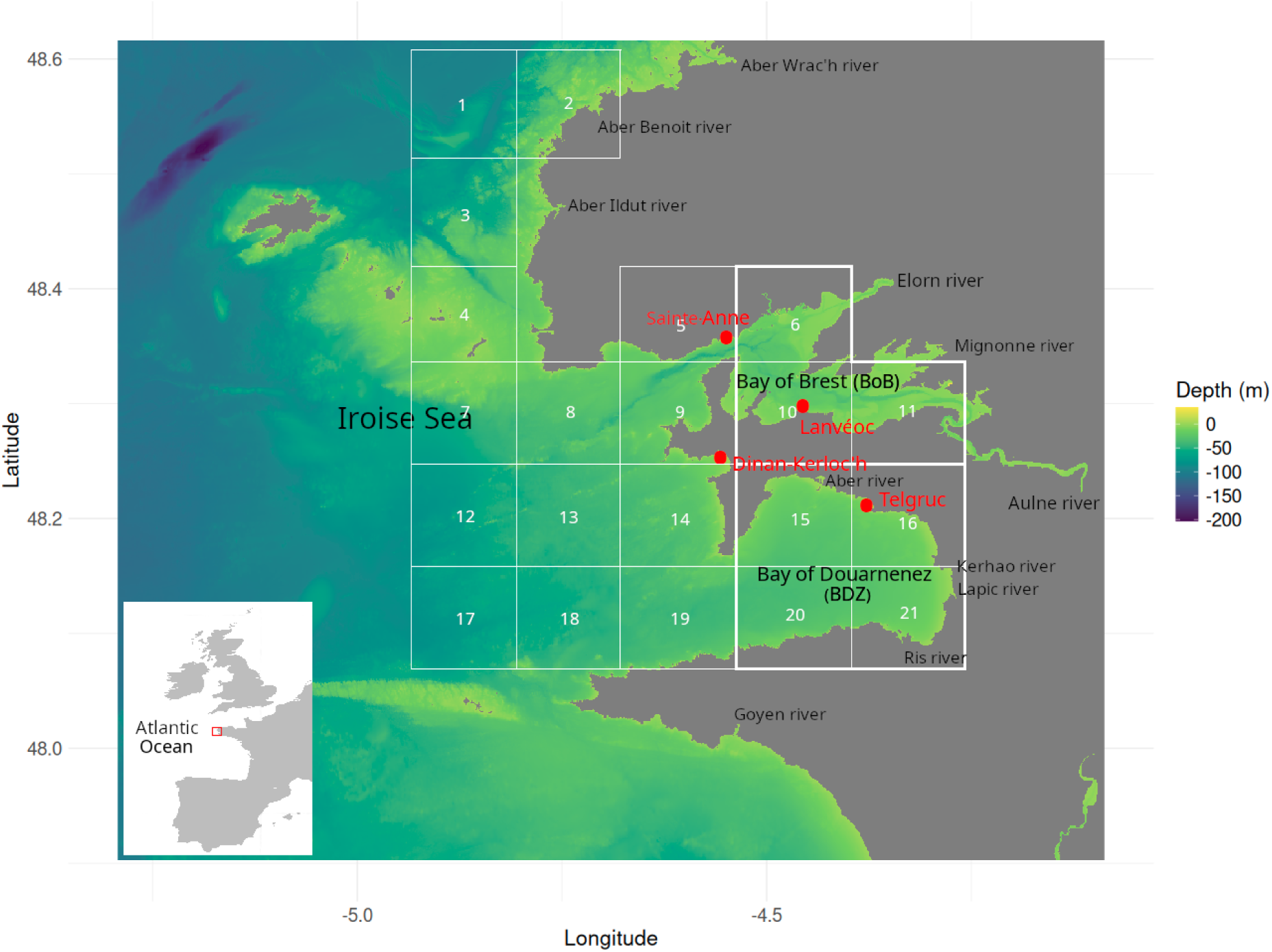
Bathymetric map of the study area (SHOM, 2015). Red dots indicate the monitoring stations in the bay of Brest (Sainte-Anne, Lanvéoc), the bay of Douarnenez (Telgruc) and the Iroise sea (Dinan-Kerloc’h). The geographical limits of this map correspond to the hydrodynamical model domain, white squares delineate the 21 zones of the Eulerian tracer.

### 2.2. Environmental parameters

During the sampling period, salinity and sea surface temperature (°C) were measured *in situ* using Cond 3110 portable thermo-salinometer (WTW, Germany). Water samples were filtered and prepared immediately after collection from shore. Nitrate, nitrite, phosphate and silicate concentrations (µmol/L) were measured by segmented flow analysis, following Aminot and Kerouel (2007). Suspended matter concentration (mg/L) was calculated following Aminot and Kérouel (2007). One liter of sea water was filtered on weighted Whatman GF/F 45 fiberglass filters (pre-combusted at 500°C for one hour) to quantify Total Suspended Sediment (TSS). The tidal amplitude (m) was recorded in Brest city harbour (048° 23’ 00.0’’ N, 004° 30’ 00.0’’ W) by the French Naval Hydrographic and Oceanographic Service (https://maree.shom.fr/). The photosynthetically active radiation (PAR, µE/m²/s) at the sea surface was calculated at each station using the Luyten et al. (1992) bulk formula which considers date, latitude, longitude, sun declination, solar radiation and fractional cloud cover (data provided by Météo-France AROME numerical model, Seity et al., 2011). The average PAR between 7am and 8pm of the four days preceding the sampling was considered as an explanatory variable of the bloom dynamics. The average PAR of the hour preceding the sampling was considered as an explanatory variable for gene expression. Daily average wave height (m) is taken from the Coriolis database observations (http://www.coriolis.eu.org). Currents were obtained using the CROCO-Iroise 3D hydrodynamic model (described in 2.8). The average speed current over the whole water column was averaged over the 12 hours preceding the sampling time, representative of a tidal cycle. Data are indicated in Supplementary material 1.

### 2.3 Cellular densities

*Pseudo-nitzschia spp.* cell abundances (cells.L^-1^) were estimated under a light microscope, using 10 mL water samples preserved in acid Lugol’s solution (2 mL.L^-1^), following the Utermöhl method. Data are indicated in Supplementary material 1.

### 2.4 Domoic acid quantification

One liter of sea water was filtered through a 47 mm glass fibre (GF/F, Whatman) to determine the intracellular toxins. The filtrates were then used for determining dissolved toxins. Samples of filters and filtrates were stored at -20°C until analysis.

To extract intracellular/particulate toxins, 250 mg glass beads (150-250 μm) and 2.5 mL MeOH/H_2_O mixture (50/50: v/v) were added to the filters and stored at -80°C for 1 hour. A mixer mill (MM400; Retsch) was used for mechanical grinding for 20 minutes at 30 Hz. The samples were centrifuged (10 min, 4,300g, 4°C), and the supernatants were filtered through a 0.2-μm nylon filter (Nanosep MF; Pall). The filtered samples were then stored at -20°C until further analysis.

To determine extracellular toxins, solid-phase extraction (SPE, Bond Elut C_18_ 200 mg cartridges; Agilent) was employed to eliminate salts and concentrate the toxins. The SPE column was conditioned with 10 mL of methanol followed by 10 mL of Milli-Q water. The sample (100 mL of seawater) was acidified at 0.2% with aqueous formic acid before passing through the SPE column. The cartridge was rinsed with 10 mL of 0.2% aqueous formic acid and dried for 1 min. Toxins adsorbed on the cartridge were eluted using 2 mL of methanol/water (50/50: v/v) into a glass vial. The eluate was stored at -20°C until analysis.

Toxins analyses were performed using Ultra Fast Liquid Chromatography (UFLC) (Shimadzu) coupled to an Sciex 4000 QTrap (triple quadrupole mass spectrometer). LC-MS/MS parameters are described in Ayache et al. (2019). A certified domoic acid standard (CNRC, Halifax, Canada) was used for external calibration range (0.3-500 ng.mL^-1^) in order to quantify domoic acid. The limit of detection (LOD) and limit of quantification (LOQ) were respectively 0.1 and 0.25 ng.mL^−1^, corresponding to 1 and 2.5 pg DA on the column. Values were analysed with Analyst 1.7.2, Sciex software. Intra and extracellular DA contents were reported in pg.mL-1. Intracellular DA was also reported in pg per *P. australis* cell. The proportion of intracellular DA was also calculated as the intracellular DA divided by the total DA.

### 2.5 RNA extraction, library preparation and sequencing

For total RNA extraction, directly (<15min) after the sampling, 4-5 L subsurface water samples were filtered on 10 µm polycarbonate filters using a peristaltic pump. Filters were frozen in liquid nitrogen in RNA later (Fisher Scientific, Illkirch, France) and stored at −80 °C until RNA extraction. Total RNA was extracted by sonicating the filters on ice (Vibra-cell 75115, Bioblock Scientific, Illkirch, France) for 30 seconds at 35% intensity in LBP buffer (Macherey-Nagel, Duren, Germany). Extraction was performed using the NucleoSpin® RNA Plus kit (Macherey-Nagel) following the manufacturer’s protocol. Library preparation was performed using the Illumina mRNA TruSeq stranded kit starting from 0.5 µg of total RNA. Paired-end sequencing was performed using 2 × 150 bp cycles on Illumina Hiseq3000 (2019 samples) and Novaseq6000 (2021 samples) at the GeT-PlaGe France Genomics sequencing platform (Toulouse, France). Generated fastQ files have been deposited to the European Nucleotide Archive (Supplementary material 1).

### 2.6 Bioinformatic analyses

Prior to read mapping, FASTQC (http://www.bioinformatics.bbsrc.ac.uk/projects/fastqc/) was used to assess read quality and quantity. TRIMMOMATIC (Version0.36) (Bolger et al., 2014) was used to trim ambiguous, low quality reads and sequencing adapters with parameters: ILLUMINACLIP:adapter.fasta:2:30:10:8 LEADING:30 TRAILING:30 MAXINFO:40:0.5 MINLEN:80.

To determine the community composition, forward trimmed reads were compared to two reference databases using blastn. The first one, the Protist Ribosomal Reference database (PR2, Guillou et al., 2012), is a manually curated database for 18S rRNA sequences, which enables complete characterization of the eukaryotic community, but at low taxonomic resolution, especially for diatoms. The second one, Diat_barcode (Rimet et al., 2019), is a manually curated database for diatom based on *rbcL*. It was used to obtain a genus level taxonomic resolution for diatoms. Reads were considered only if their minimum e-values were <10⁻³⁰ against Diat_barcode and <10⁻⁷⁰ against PR2. For each retained read, if all database sequences with the minimum e-value belonged to the same phylum (PR2) or genus (Diat_barcode), the read was assigned to that phylum or genus. All other reads were not considered. Relative community composition was obtained by dividing the number of reads assigned to a given taxonomic level (phylum or genus) by the total number of taxonomically assigned reads in each sample. Only phylum or genus with a relative abundance >5% were considered.

To determine *P. australis* gene expression dynamics, BWA-MEM (Li et al., 2013), was used to align all trimmed reads to a metareference corresponding to the combination of 315 species specific reference transcriptomes, representing 213 unique genera (Metegnier et al., 2020; Prigent et al. 2025). It mostly corresponded to the resources developed during the Marine Microbial Eukaryotic Transcriptome Sequencing Project (MMETSP, Keeling et al., 2014) with the addition of reference transcriptomes obtained for three *Pseudo-nitzschia* species (*P. australis*, *P. fraudulenta* and *P. pungens*) based on local strains (Lema et al., 2019). In addition to these three species, reference transcriptomes of *P. arenysensis*, *P. delicatissima* and *P. heimii* were also present in the metareference. Samtools was used to discard reads displaying low quality alignments (MapQ<10), to remove read pairs that did not align to the same transcript and to generate the raw read count expression matrix (Li et al., 2009).

In each sample, the proportion of *P. australis* among *Pseudo-nitzschia* species was obtained by dividing the number of reads assigned to *P. australis* by the sum of reads assigned to all *Pseudo-nitzschia* species. This proportion was multiplied by the *Pseudo-nitzschia spp.* abundances, quantified using optic microscopy, to estimate *P. australis* absolute abundances.

For gene expression analysis, samples with less than 500,000 reads aligning to *P. australis* transcripts were discarded. Transcripts displaying a read coverage < 5 in more than half of the samples were discarded. Following these filtering steps, a total of 24 samples and 11,924 transcripts were analyzed. Deseq2 Variance Stabilizing Transformation was used to normalize read counts (Love et al., 2014). Transcripts with similar expression dynamics were grouped into modules of co-expressed transcripts using WGCNA (Langfelder and Horvath, 2008) with a soft-thresholding power of 9 and bicor correlation. Module identification was performed using dynamic tree cut with minimum cluster size of 350 transcripts, with deepSplit=2 and cutHeight=0.98. The package WGCNA was used to extract an eigengene representing the expression dynamic of each module.

The *P. australis* reference transcriptome was annotated using eggNOG-mapper 2.1.9 (Cantalapiedra et al. 2021) using Diamond (Buchkink et al. 2021) on more-sensitive mode against the Uniprot/Swissprot database with a maximum evalue of 10^-3^ and considering KEGG annotations. For each expression module, Kegg pathway enrichment was tested using Fisher Exact tests followed by FDR correction with a q-value threshold set at 0.05. Fisher Exact tests odds ratios (ORs) were reported. Only KEGG categories represented by at least five expressed transcripts were considered.

### 2.7 Statistical analysis

All statistical analyses and graphics were performed with R 4.2.2. Three distinct datasets were considered. The first one corresponded to the environmental parameters and included nutrient concentrations, temperature, salinity, PAR, current, TSS and wave height. The second one corresponded to the *Pseudo-nitzschia* dataset and included *Pseudo-nitzschia spp.* abundances, *P. australis* abundances, extra and intracellular domoic acid concentrations. The third one corresponded to the eigenene expression for each gene module.

Relationships between the centered and scaled 1. *Pseudo-nitzschia* dataset as response variables and environmental dataset as explanatory variables and 2. eigengenes expression as response variables and the two other datasets as explanatory variables were explored using two Redundancy Analysis (RDA, vegan package; Oksanen et al., 2022). A total of 39 and 24 samples, respectively, had no missing data and were included in the two RDAs. Dates (as a continuous variable) and sampling sites (dummy-coded, four sampling sites coded in the first RDA, two sampling sites (Telgruc and Others) coded in the second RDA) were included as additional explanatory variables. Correlation among explanatory variables (excluding sampling sites) was first assessed by pairwise Spearman correlations; when rho > 0.7, a single variable was kept. In the first RDA Silicate, Phosphate and Salinity were not considered as they correlated with Nitrate+Nitrite. In the second RDA: 1. Phosphate, the proportion of intracellular domoic acid and the quantity of extracellular domoic acid were correlated with the sampling dates; 2. Salinity and Silicate correlated with Nitrate+Nitrite. In the two RDAs, *Pseudo-nitzschia spp.* density and intracellular domoic acid concentrations (pg.mL-1) were excluded as they correlated with *P. australis* density. Remaining environmental variables were further screened using Variance Inflation Factors (VIF, vif.cca() function) on the full model, removing variables iteratively starting from the highest VIF until all VIF < 6. This led to the exclusion of Current in the first RDA. Forward selection of explanatory variables was performed using the forward.sel() function (adespatial package; Dray et al., 2023) with a double stopping criterion based on permutation tests (α = 0.05) and the adjusted R² of the full model. The significance of the final model and marginal effects of individual terms were tested using permutation-based ANOVA (999 permutations). Adjusted R² was calculated to assess the proportion of variance explained.

### 2.8 Hydrodynamical model

Hydrodynamic simulations were performed using the CROCO model (Coastal and Regional Ocean Community model), derived from the Regional Oceanic Modeling System (ROMS) (Shchepetkin and McWilliams, 2005). This three-dimensional free-surface model solves the simplified primitive Navier-Stokes equations, under the assumptions of hydrostaticity, Boussinesq, and incompressibility, as well as the nonlinear advection-diffusion equation for tracers such as temperature, salinity, or any other possible conservative or non-conservative tracer. The model also takes into account a wetting and drying capability for shallowest areas, due to the tide. A detailed description of the CROCO model and its Fortran code is available on the open repository Auclair et al. 2022 (https://zenodo.org/records/17642275).

Horizontally, the model spreads on a 404 x 441 square cells grid with a spatial resolution of 200 meters (the total model extent is presented on Fig. 1). Vertically, 30 sigma levels are considered with no refinement at the surface and bottom since the model is predominantly coastal and depths do not exceed 200 m. The bathymetry of the model (Fig. 1) was built from the bathymetric DEM dataset of the Atlantic coast with a resolution of 0.001° (∼ 111 meters) (SHOM, 2015).

The daily flows of the 7 largest coastal rivers within our area (Aulne, Elorn, Mignonne, Aber Ildut, Aber Benoît, and Aber Wrac’h) are taken from the HYDRO database (https://www.hydro.eaufrance.fr). For the smaller rivers (Sainte-Anne, Aber, Kerharo, Lapic, Ris), which are not included in the HYDRO database, monthly or annual flow climatologies were created based on literature data when available or on catchment area surface data.

The atmospheric forcings are derived from the AROME model (Météo France, Seity et al., 2011). At the three open boundaries of the model (North, South and West), the following boundary conditions were applied :

- Water level variations due to tides are based on the Previmer Atlas with 37 tidal waves (Pineau-Guillou, 2013).
- Temperature and salinity tracers are based on the outputs of a larger model (MARS3D-MANGAE2500 configuration, http://marc.ifremer.fr/) are used after interpolation on the model grid.

The initial conditions for the two tracers are derived from the MARS3D-MANGAE2500 outputs interpolated on the grid. In order to move away from these initial conditions and to obtain a coherent signal for these parameters, a 1-month spin-up was systematically performed before each simulation.

#### Validation

In order to assess the model’s ability to reproduce real hydrodynamic conditions in the area, simulations were compared to *in situ* measurements. Validation focused on the following variables: the simulated water height (tide) was compared to 3 tide gauge stations : Brest (48.3829°N; -4.4950°E), Le Conquet (48.3591°N; -4.7808°E) and Audierne port (48.0216°N; -4.5375°E); simulated baroclinic current speeds and directions were compared with ADCP acoustic profiler datasets (Petton et al., 2016) placed in the Anse de l’Auberlac’h (48.3272°N, -4.4297°E), the Anse du Fret (48.2916°N; -4.4841°E), the Lomergat (48.2877°N; -4.3696°E) and the Anse du Roz (48.3207°N; -4.3399°E), and simulated temperature and salinity were compared to data coming from two high-frequency monitoring stations, the COAST-HF MAREL-Iroise buoy (48.357°N; -4.582°E, Rimmelin-Maury et al., 2023), and the Pointe du Château station (48.335°N, -4.319°E, Petton et al., 2022).

Detailed validation results are presented in Supplementary material 2 but, in summary, the simulated tide is good during all the study period. With spring tide amplitudes ranging between 4 to 7.5 m, maximum RMSD was 30 cm at the southernmost station Audierne and around 20 cm at other stations. Simulated current speeds are also satisfactory with mean RMSD and MAE of 0.06 m/s and 0.04 m/s respectively. Finally, the simulated salinity and water temperature also show good results even if the model tends to over-estimate the salinity at station Sainte-Anne with MAE around 0.7 PSU. Overall, the validation exercise, although incomplete (carried out only at few stations, and for a limited time period), proved satisfactory to confidently address the movement of tracers in the study of connectivity.

#### Connectivity

In order to investigate the water masses movements between different areas in this study site, our hydrodynamical model was used to simulate the transport of conservative Eulerian tracers, initialized in 21 different zones (Fig. 1). Each zone (50 x 50 model grid cells, approx. 10x10 km), represents both a starting point for a conservative Eulerian tracer and an observation zone for the arrival of tracers originating from the other 20 zones.

All tracers were initialized on the 26th of February 2021 with a theoretical concentration of 1 and the simulations run during 1 month. The initial mass of each tracer obviously depends on the local bathymetry thus, in order to calculate comparable indicators, the initial mass of each tracer at the beginning of the simulation was calculated in the starting area, and all tracer quantities reaching the ending areas were expressed as a percentage of this initial mass. Two connectivity indicators were calculated : the maximum percentage of initial tracer mass (Pmax) reached during the simulation, and the time associated with this maximum (Tmax).

#### Local e-flushing time and dilution rates

The hydrodynamical flushing that experience any developing plankton cells in a particular area, has been assessed using the e-flushing time (expressed in day) defined by Koutitonsky et al. (2004) : it is the time needed for the dilution to drop definitively the tracer below 37% (*i. e.* e^-1^) of its initial mass. Considering that the water renewal, due to waters coming from outside the area, is quasi-exponential, the dilution during the first day can be calculated as following : *d* =− *ln*(*B*/*B*_0_) with *B,* the remaining mass of tracer after 1 day simulation and *B_0_*, the initial mass of tracer. For those indicators, 15 shorter simulations were carried out (duration 10 days), with re-initialization of tracers at each sampling date from the 26^th^ of February to the 2^nd^ of April 2021.

### 2.9 Modeling bloom dynamics

Temporal dynamics of *Pseudo-nitzschia australis* abundance at each station were modelled using Generalized Additive Models (GAM) with a Gamma distribution and log link function. Abundance was modelled as a smooth function of time (days), using cubic spline basis functions. The degree of smoothness was evaluated by testing values of k ranging from 3 to 6. All models were fitted using the mgcv package in R. The fitted models were then used to estimate time lags between stations in exceeding a threshold of 10,000 cells L⁻¹.

To determine if the difference in dilution between the Bay of Brest and the Bay of Douarnenez alone may explain the observed time lag between stations, simple exponential growth models with dilution were considered as N(t)=N_0_e^(μ−D)t^ where N_0_ and N(t) are the cell densities (cell.l^-1^) at time 0, µ and D are the division and dilution rates (d^-1^) and t the time (d). Division rates ranging from 0.4 to 1 were considered to determine if they were compatible with observed time lags.

## 3. Results

### 3.1 Environmental conditions

Overall, the seasonal transition from winter to spring was the strongest environmental signal recorded during the study period at each four stations (Figure 2). This was especially visible in terms of nutrient concentrations, with inorganic nitrogen, silicate and phosphate concentrations decreasing from an average of 27µM, 15µM and 0.36µM on February 26^th^ down to 7 µM, 3 µM, and 0.07µM on April 16^th^, respectively (excluding Sainte-Anne station). The winter-spring transition was also visible in terms of PAR and salinity increase, but also wave height and total suspended solid decrease toward the end of the sampling period. Temperature tended to increase in the two stations of the Bay of Brest while in the other two there was an initial increase and a drop in temperature toward the end of the survey. Although there is no temporal trend in terms of current, the current speed was systematically higher in the two stations from the bay of Brest (even higher at the Sainte-Anne station). The Sainte-Anne station stands out in terms of environmental conditions and especially nutrient concentrations, with strong changes from one day to the next and extremely high transient concentrations. For instance, inorganic nitrogen exceeded 50 µM seven times at Saint-Anne with a maximum concentration of 285µM, while the inorganic nitrogen concentrations were always below 50 µM at the other stations. At the Telgruc station, there was a drop in phosphate and silicate concentrations between March 19^th^ and 30^th^ that led to a strong increase in N:P and N:Si ratio.

**Figure 2:**
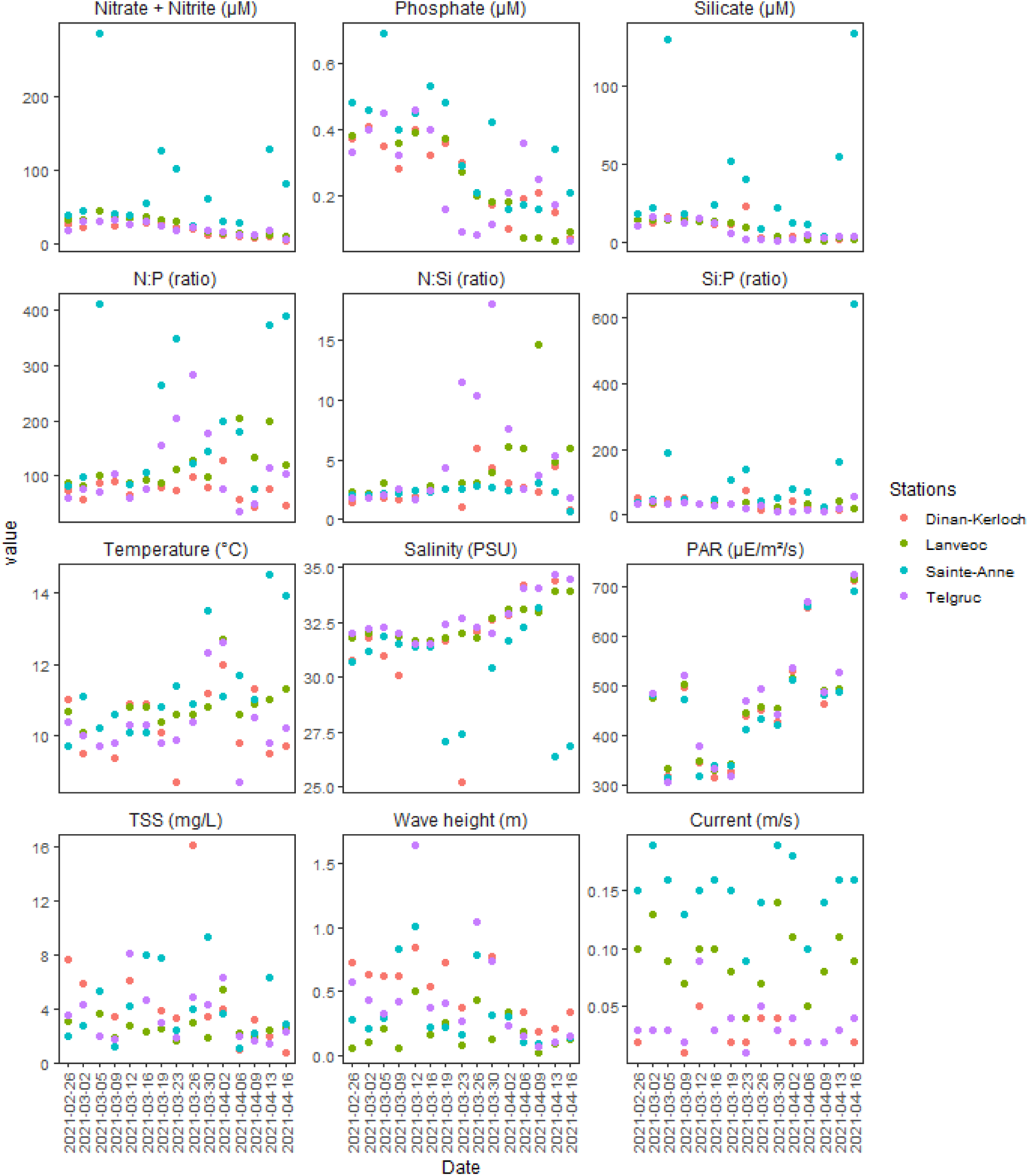
Environmental conditions. Evolution of 12 abiotic parameters as a function of time at the four sampled stations. Parameters and units of the y axes are indicated above the graph.TSS stands for total suspended sediment. Colours indicate the four stations.

### 3.2 Community composition

Bacillariophyta, i.e. diatoms, tend to dominate the community throughout the survey (Average relative abundance of 51%). Relative abundances were slightly lower in the two stations of the Bay of Brest (average of 48%) compared to Dinan-Kerloch and Telgruc (average of 55%). There were punctual drops in diatom relative abundances, especially between March 26^th^ and April 2^d^ at Telgruc (Figure 3A).

**Figure 3:**
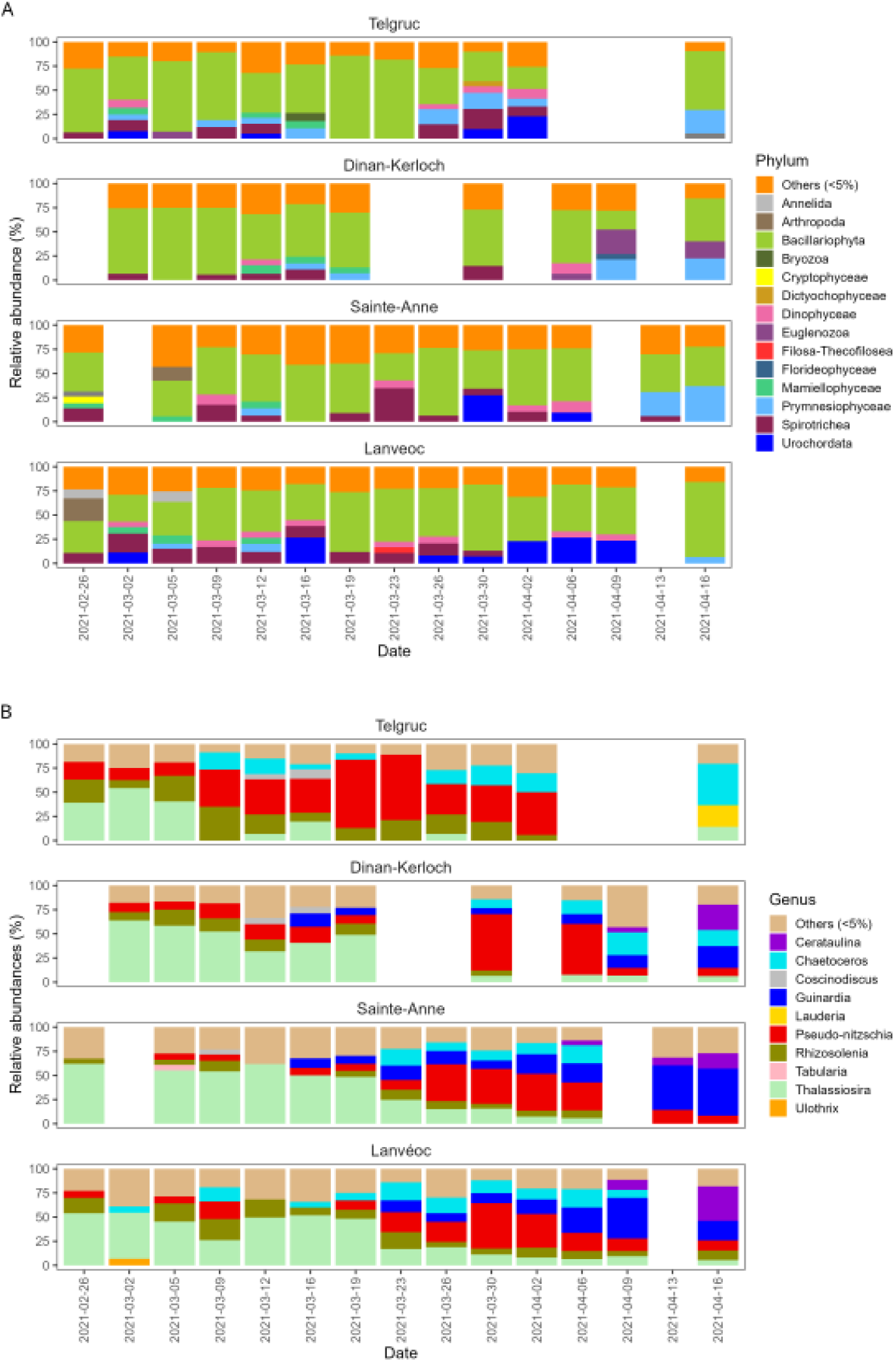
Relative abundances of Phylum (A) and Diatom genera (B) at the four stations through time. Phylum and diatom relative abundances are based on the alignment on the PR2 (18S rRNA) and Diat_barcode (rbcL) databases, respectively. Missing samples are in white. Only taxa with relative abundance > 5% are indicated.

Within the diatoms (Figure 3B), 10 genera represented more than 5% in at least one sample. Members of the *Thalassiosira* genus represented a major fraction of the diatom community at the beginning of the survey, often representing more than 50%. *Thalassiosira* relative abundance quickly dropped at the Telgruc station, while relative abundance of *Pseudo-nitzschia*, and to a lesser extent, of *Chaetoceros* increased. The same transition from *Thalassiosira* to *Pseudo-nitzschia* (and *Chaetoceros*) also occurred at the other stations, but much later. For instance *Pseudo-nitzschia* increased from 14 to 39% of relative abundance on March 9^th^ at Telgruc, while comparable increases occurred at least 14 days later at the other stations (from march 23^d^ to 30^th^). Deviation from this spatio-temporal shift in community composition were also observed. For instance, the increase in *Pseudo-nitzschia* abundance co-occurred with an increase in *Guinardia* in all stations but Telgruc. Moreover, *Rhisosolenia* relative abundance remained somehow similar during the spatio-temporal shift and, toward the end of the survey, Sainte-Anne and Lanveoc communities were dominated by *Guinardia* and *Cerataulina,* the same genera were associated with *Chaetoceros* at Dinan-Kerloch, while at Telgruc *Chaetoceros* and *Lauderia* were dominant.

### 3.3 *Pseudo-nitzschia australis* abundances and domoic acid concentrations

The bloom dynamic differed across time and stations (Figure 4A and B; RDA F=6.0, p-value=0.001, permutation-based ANOVA, adjusted R²=0.21). More specifically, the bloom occurred earlier in Telgruc (Telgruc effect, F=5.3 p-value=0.002; Dates effect, F=6.7 p-value=0.004). Absolute abundances of *P. australis* initially increased in Telgruc, with densities between 40,000 and 400,000 cells per liter between March 9^th^ and 23^d^ (Figure 4A). At the other stations, densities were more moderate, reaching maximum values between 35,000 and 55,000 cells per liter on March 26^th^ and 30^th^. GAMs (k from 3 to 6) were fitted to the *P. australis* densities of the four sampling sites (Figure 4A). Based on these models, the time lags to reach 10,000 cells/L at the various sampling sites are 9 to 13 days between Telgruc and Dinan-Kerloch, 24 to 28 days between Telgruc and Lanveoc and 24-25 days between Telgruc and Sainte-Anne. Moreover, at Telgruc, the proportion of intracellular domoic acid was between 52% and 82% from February 26^th^ to March 23^d^ before dropping quickly. In contrast, at the other stations, the proportion of intracellular domoic acid remained below 30% between March 19^th^ and 26^th^, when *P. australis* abundances were increasing at these stations (Figure 4A). At Telgruc, cell densities dropped after March 23^d^ (Figure 4A). This drop was concomitant with a decrease in the proportion of intracellular domoic acid (Figure 4A) and was preceded on March 19^th^ by a strong decrease in phosphate concentration that strongly affected nutrient ratio (Figure 3A). Overall, there was a tendency for extracellular domoic acid to increase with time (Figure 4A). Finally, intracellular domoic acid content averaged 13.4±7.5 pg/cell (mean±sem) and was not strongly correlated with environmental variables or sites (Figure 4A).

**Figure 4:**
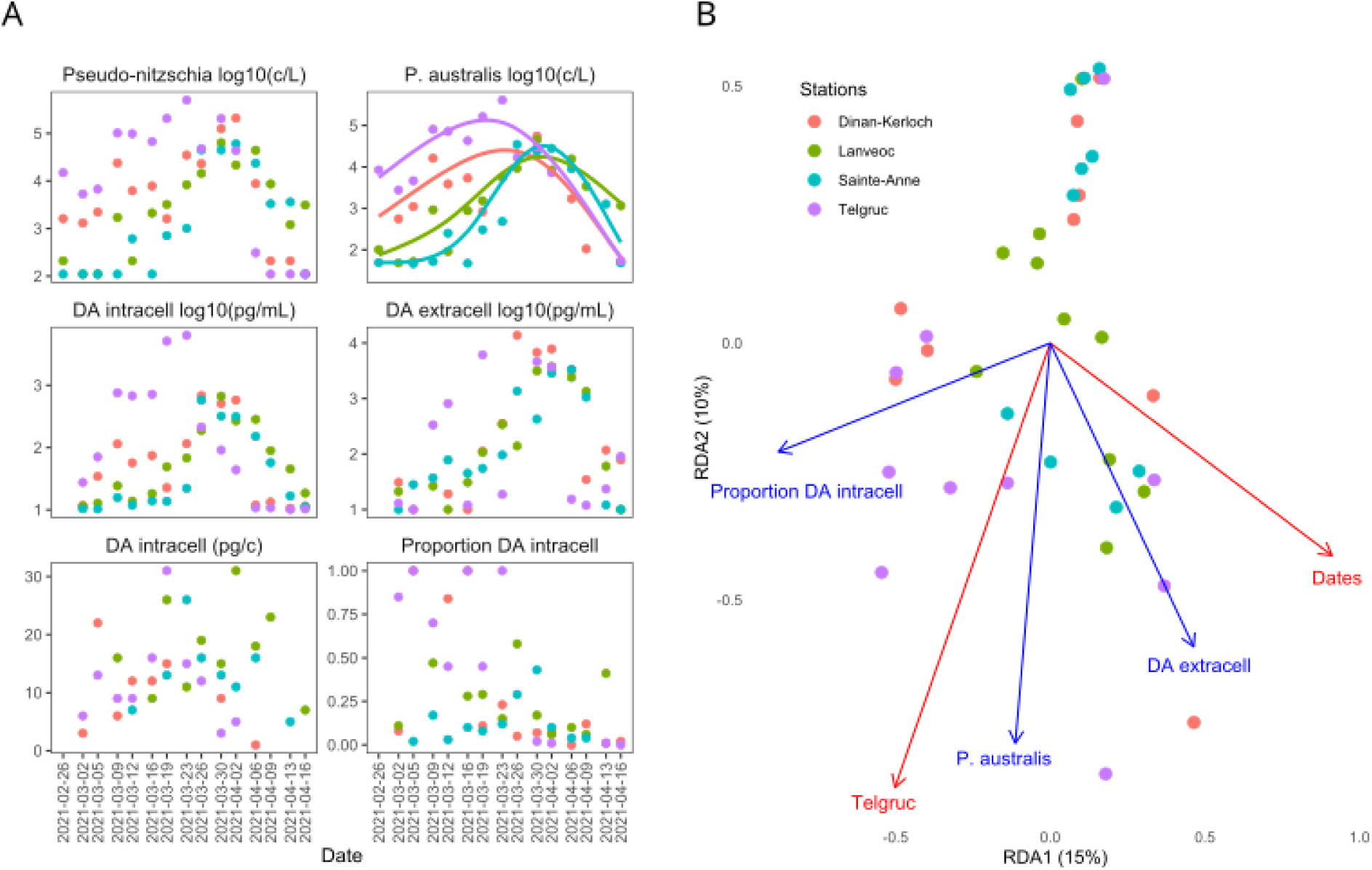
Pseudo-nitzschia australis bloom dynamics. Pseudo-nitzschia abundances and domoic acid (A) and RDA analysis indicating significant relationships with environmental parameters, dates and locality (B). Gam-smoothed (k=4) bloom curves are fitted to P. australis abundances for each of the four stations. Each dot corresponds to a sample, with colors indicating the sampled stations. Blue arrows indicate the Pseudo-nitzschia parameters used in the RDA analysis. Red arrows indicate the significant explanatory variables.

### 3.4 Pseudo-nitzschia australis gene expression

*Pseudo-nitzschia australis* expression was analyzed for 11,924 transcripts in 24 samples. A total of 5 expression modules regrouping transcripts displaying similar expression dynamics across samples were identified (Figure 5A). Gene expression was correlated with time (Figure 5B; RDA, time effect, F=7.8, p-value=0.001, permutation-based ANOVA), *P. australis* abundance (Figure 5B; RDA, *P. australis* effect, F=6.7, p-value=0.001, permutation-based ANOVA), as well as with the average PAR of the hour preceding the sampling (Figure 5B; RDA, PAR effect, F=4.0, p-value=0.008; full model, F=6.1, p-value=0.001, permutation-based ANOVA, adjusted R²=0.40).

**Figure 5:**
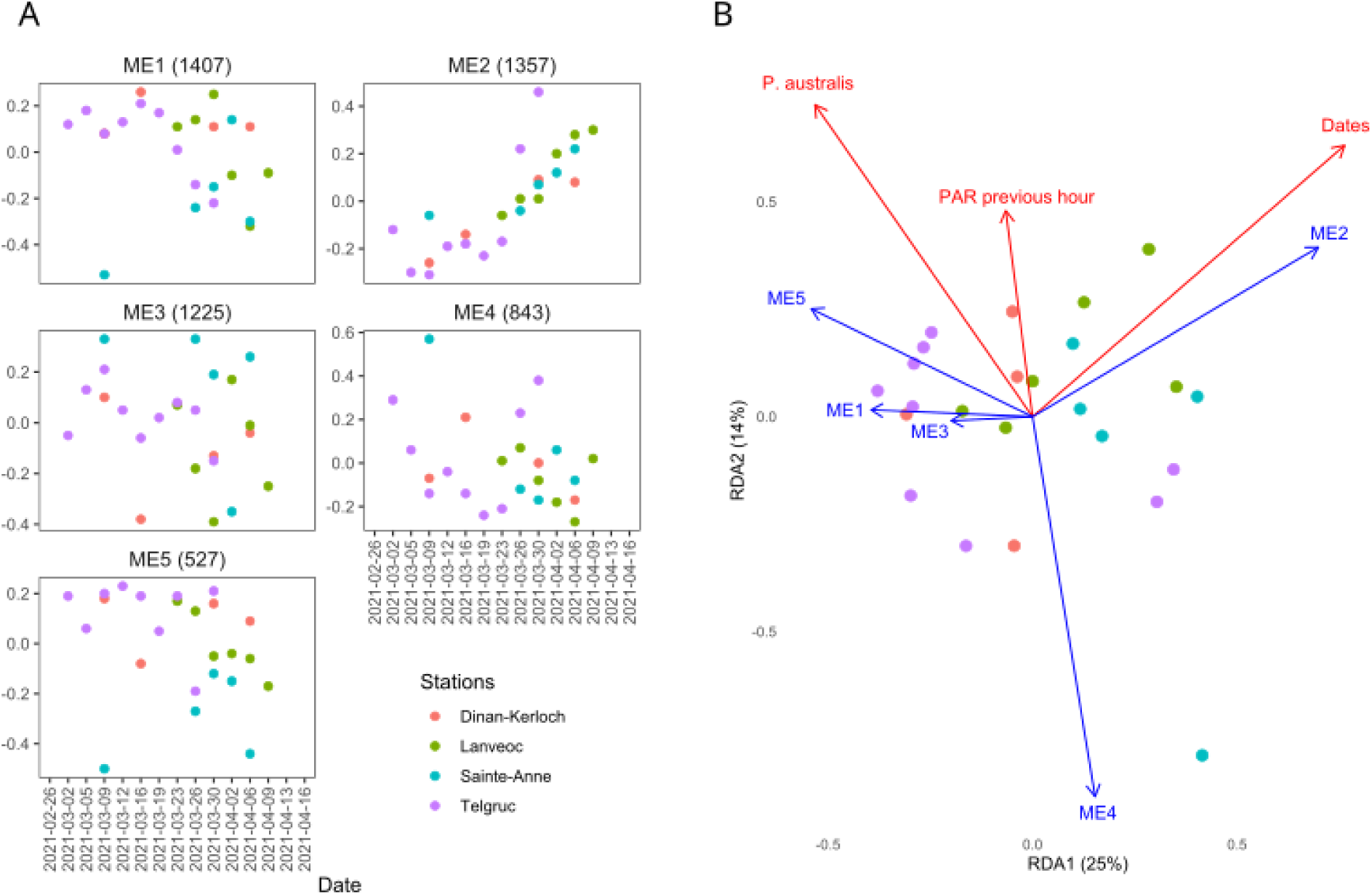
Pseudo-nitzschia australis gene expression. Eigengene expression for each gene expression module (ME1 to 5; A) and RDA analysis indicating significant explanatory variables (B). Each dot corresponds to a sample, with colors indicating the sampled stations. Blue arrows indicate the eigengenes expression used in the RDA analysis. Red arrows indicate the significant explanatory variables.

More specifically; the transcripts of the first expression module (ME1, 1,407 transcripts) showed high expression up to the maximum abundance before decreasing sharply as abundances declined. This was especially noticeable at Telgruc and Lanvéoc (Figure 5A). Transcripts associated with ribosomes and involved in various metabolic processes such as carbon fixation and metabolism, amino acid, fatty acid and secondary metabolites metabolism tend to be over-represented in this module (Figure 6). The expression of the 1,357 transcripts of the second module (ME2) displayed a strong increase with time. More specifically, very low expression levels were observed in Telgruc until the maximum abundance of *P. australis*, followed by a very sharp increase. At the other stations, ME2 expression also increased with time, but moderate expression levels were observed before the maximum abundances. Transcripts related to cellular junctions as well as apoptosis and necroptosis were over-represented in this module (Figure 6). Visual inspection of the transcripts related to these two later functions indicated that they did not correspond to specific diatom programmed cell death (PCD) components such as metacaspases (see discussion). Instead, it was characterized by broader functions associated with cytoskeletal dynamics, intracellular trafficking, proteostasis, energy metabolism and cellular stress (Supplementary material 3). Transcripts of the third module (ME3, 1,225 transcripts) were not strongly correlated with bloom or environmental parameters (Figure 5A, B). A few functions related to protein turnover were over-represented in this module. The fourth module (ME4, 843 transcripts) displayed strong negative correlation with the average PAR of the hour preceding the sampling (Figure 5B). It included an over-representation of genes related to apoptosis and phagosomes (Figure 6). Finally, expression of the fifth module (ME5, 527) decreased with time but did not display any functional enrichment (Figure 5, 6).

**Figure 6:**
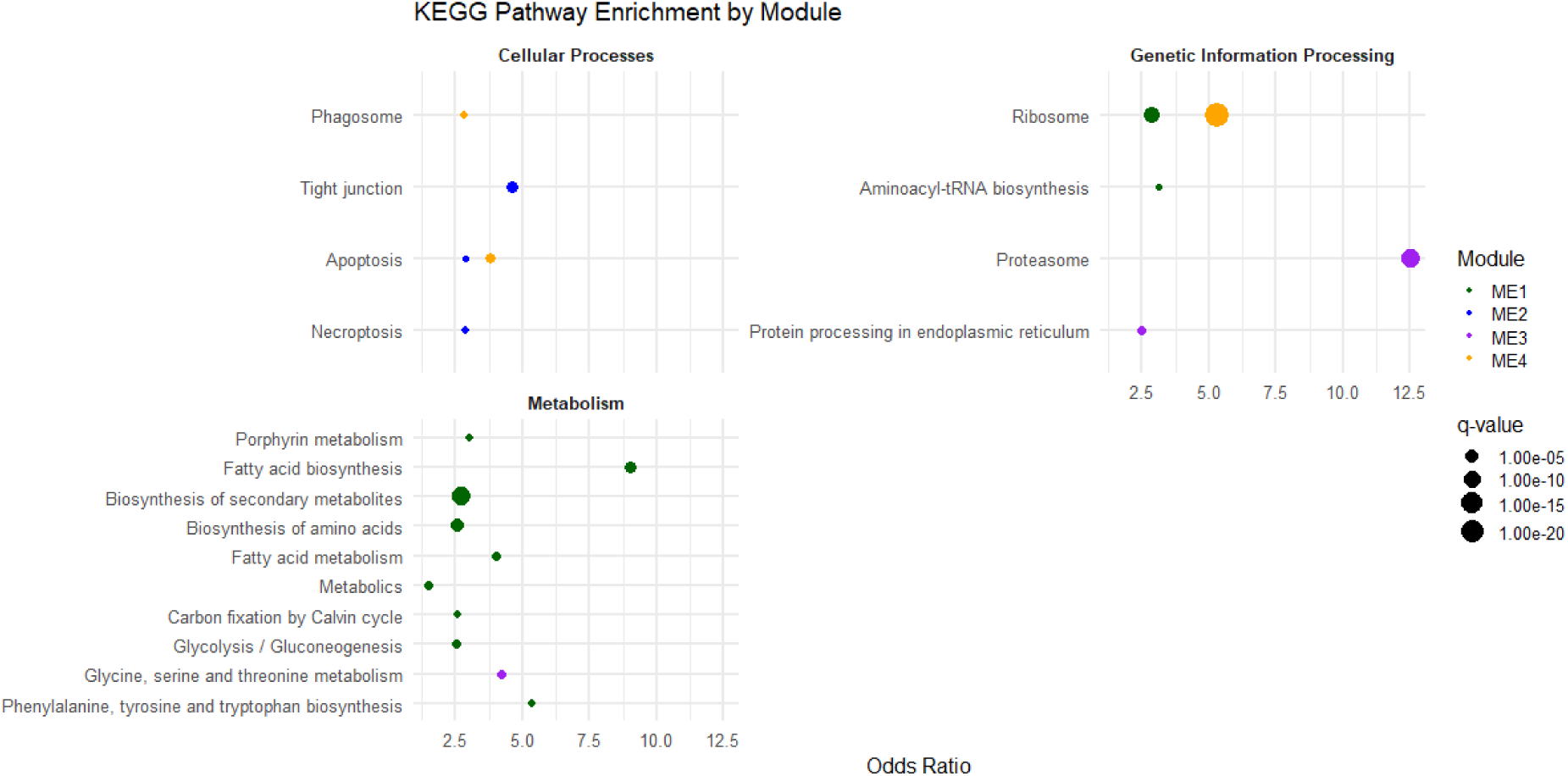
Expression modules KEGG pathway enrichment. The odds-ratio of the significant KEGG pathways are indicated, with the size of the circles corresponding to the q-values. Pathways are grouped into three broad categories related to Cellular processes, Genetic information processing and Metabolism. Colors indicate the expression modules.

### 3.6 Hydrodynamic modelling

The results presented above indicated that the *P. australis* bloom started, peaked with higher densities and decreased earlier in the Telgruc station than in the three other stations. In the following section, two hypotheses were tested regarding the putative influence of the hydrodynamism on this time lag. The first one is that bloom onset was truly anticipated in Telgruc and that hydrodynamic connectivity seeded the bloom at the other stations. The second hypothesis is that stronger hydrodynamic dilution in the Bay of Brest compared to the Bay of Douarnenez may result in cell densities increasing earlier and reaching higher levels even if growth rates are similar in the two bays. To test these hypotheses, three dimensional hydrodynamic simulations were performed.

#### Connectivity in the Iroise Sea

The Tmax and Pmax connectivity matrices are presented in Supplementary material 4. All 21 zones are connected with each other at the end of a 1 month simulation. Nonetheless, differences appear in the maximum percentage of the different tracers reaching a particular zone (Pmax) and the times needed to reach those maximums (Tmax). In the following we will consider that the higher the Pmax (respectively the lower the Tmax), the higher the connectivity between two zones. Our results show that during our study period (end of February 2021 to the beginning to April 2021) tracers coming from the bay of Douarnenez (hereafter BDZ) always reach the bay of Brest (hereafter BoB) with higher Pmax and lower Tmax than the contrary, which indicates a stronger South-to-North connectivity. Indeed the tracers coming from the BoB (released in zones 6, 10 and 11) tend to be more directed towards the West and North-West (*i. e.* towards the Iroise Sea) than to the south towards the BDZ. If we simplify the analysis of connectivity matrices, by grouping the different zones into three large areas (BDZ: zones 15, 16, 20 and 21, BoB : zones 6, 10 and 11, and the Iroise Sea: zones 4, 7, 12 and 17), the mean Pmax/Tmax is 0.9%/29.6d for BDZ towards BoB, whereas it is 0.5%/22.5d for BoB towards BDZ, meaning that tracers arrive faster but more diluted when heading South than when heading North (Figure 7). When looking at the connectivity between the two bays and the Iroise Sea, it appears that the BoB is more connected to the Iroise Sea than BDZ (mean Pmax/Tmax is 5.2%/8.4d for BoB towards the Iroise Sea and 3.3%/16.3d for BDZ towards the Iroise Sea). The Tmax between BDZ and BoB were between 28 and 31 days, which was extremely close to the time lags to reach 10,000 cells/L at the sampling sites situated in BDZ and BoB (24-28 days, Figure 2A).

**Fig. 7.**
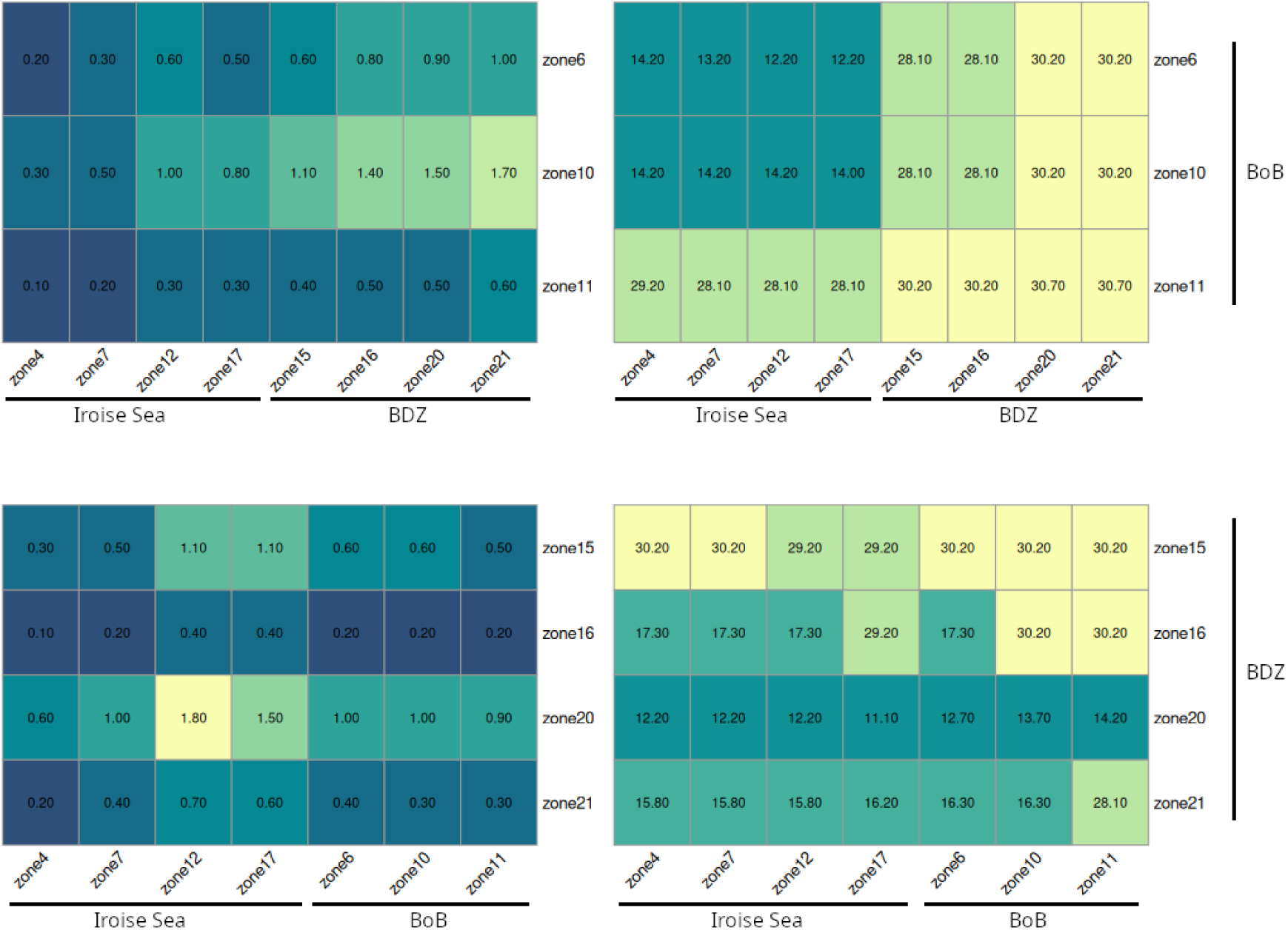
Simplified connectivity matrices presenting Pmax (left panels, expressed in % of the initial tracer mass) and Tmax (right panels, expressed in days) results. The departure areas are presented on the bottom of each matrix, and the arrival areas are presented on the right. For example, the top panels present the Pmax and Tmax of tracers coming from the BDZ and the Iroise Sea and arriving in the BoB.

#### Local e-flushing times and dilution rates

The table 1 presents the calculated e-flushing times and one-day dilution rates experienced by the two bays during all the study period. For the sake of clarity the tracers initiated in areas 6, 10 and 11 (respectively areas 15, 16, 20 and 21) were pooled and the results presented as BoB (respectively BDZ).

**Table 1.** Local e-flushing times and one-day dilution rates calculated with the hydrodynamical model. LT and HT stand respectively for low tide and high tide and give the moment of tracer release relative to the tide.

| Date of tracer initialization | Tide (Coef) | 1st day dilutions (d<sup>-1</sup>) |  | e-flushing times (d) |  |
| --- | --- | --- | --- | --- | --- |
|  |  | BoB | BDZ | BoB | BDZ |
| 21/02/26 00:00:00 | LT+2h (79) | 0.40 | 0.13 | 4.8 | 6.9 |
| 21/02/28 00:00:00 | LT (101) | 0.50 | 0.12 | 5.9 | >9.0 |
| 21/03/02 00:00:00 | LT-1h (105) | 0.41 | 0.22 | >9.0 | >9.0 |
| 21/03/05 00:00:00 | LT-3h (71) | 0.26 | 0.10 | >9.0 | >9.0 |
| 21/03/09 00:00:00 | LT+4h (48) | 0.18 | 0.06 | 8.9 | >9.0 |
| 21/03/12 00:00:00 | LT+2h (83) | 0.35 | 0.18 | 9.4 | >9.0 |
| 21/03/16 00:00:00 | LT-1h (88) | 0.35 | 0.08 | >9.0 | >9.0 |
| 21/03/19 00:00:00 | LT-2h (61) | 0.26 | 0.10 | >9.0 | >9.0 |
| 21/03/23 00:00:00 | HT (28) | 0.10 | 0.03 | 8.3 | >9.0 |
| 21/03/26 00:00:00 | LT+3h (65) | 0.29 | 0.07 | 5.4 | >9.0 |
| 21/03/30 00:00:00 | LT-1h (112) | 0.50 | 0.08 | 8.4 | >9.0 |
| 21/04/02 00:00:00 | LT-3h (88) | 0.30 | 0.13 | >9.0 | >9.0 |

The simulations highlight the differences in hydrodynamic behaviour of the two bays. Throughout the study period, the Bay of Douarnenez exhibits higher e-flushing times and lower dilution rates than the Rade de Brest. Indeed, short term dilution rates that prevail in the BoB are on average 3.3 times higher than in the BDZ. Moreover, in the BDZ, the e-flushing time was minimum at the beginning of the study period (6.9 days the 2021-02-26) and later remains always higher than 9 days while it was frequently below 9 days in the BoB (the minimum value, 4.8 days, was also calculated at the beginning of the time series). This brings out the more energetic nature of the BoB’s hydrodynamics compared to that of the BDZ, which appears to be a more quiet and confined area. Interestingly, the correlation between the tidal coefficients and the dilution rates is much more pronounced for the BoB (R^2^=0.88, overall linear model p-value < 1e^-6^) than for the BDZ (R^2^=0.33, overall linear model p-value = 0.03). This implies that the dilution in the BoB is mainly due to the tidal currents while it is less the case in the BDZ with wind-driven currents probably having a higher impact on the dilution.

In order to determine if the difference in dilution between the two bays may be a potential explanation for the observed time lag between *P. australis* accumulation in BDZ and BoB, simple theoretical exponential growth models with dilution were considered. Considering the average modelled dilution rate of 0.11 d^-1^ in BDZ and 0.33 d^-1^ in BoB, a growth rate ∼0.45 d^-1^, mimicked the observed 24 to 28 days lag (Figure 4A) to go from 100 cell.l^-1^ to 10,000 cell.l^-1^ in the two bays.

**Figure 8:**
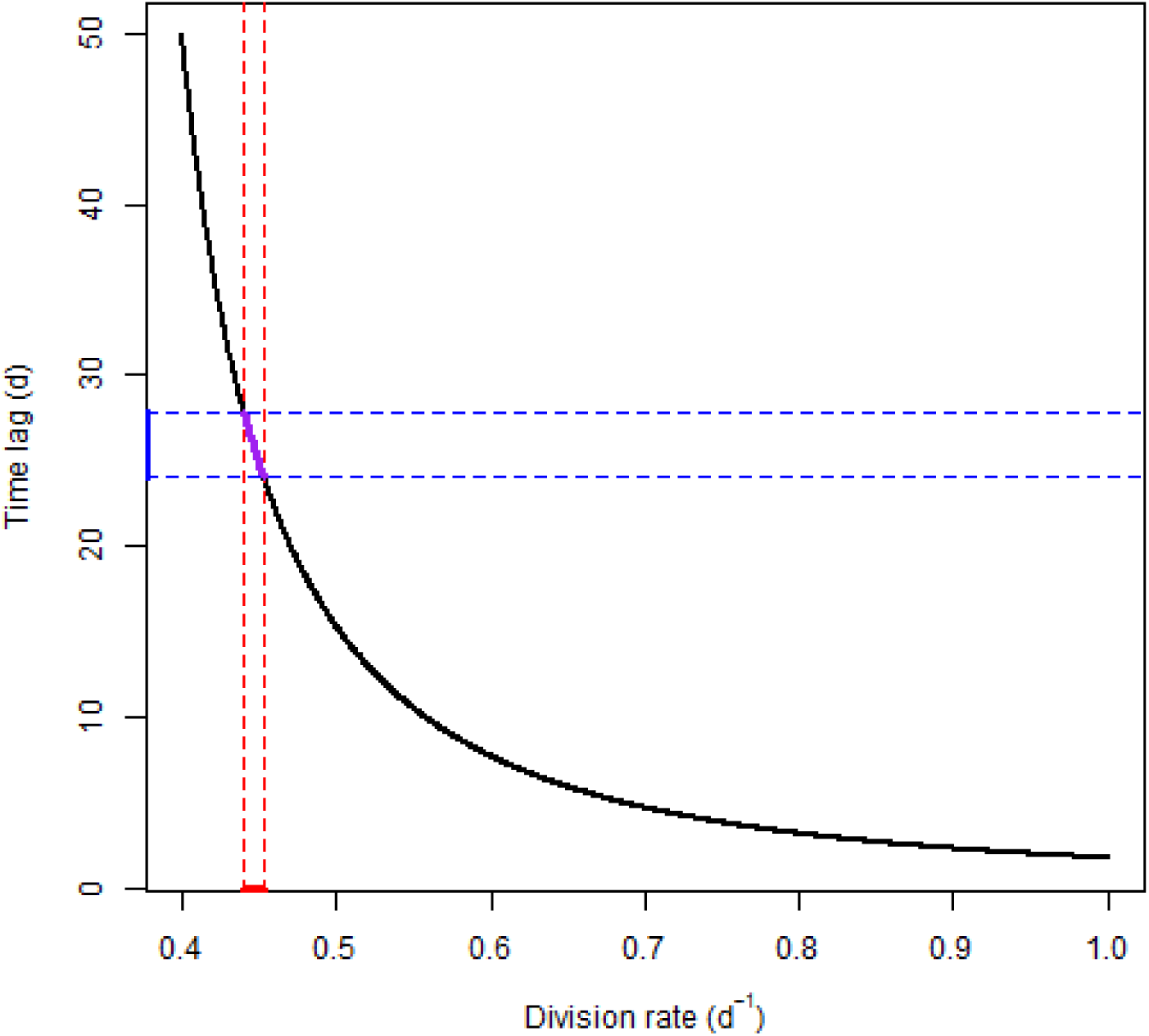
Time lag (days) to go from 100 cell.l^-1^ to 10,000 cell.l^-1^ with a dilution rate of 0.11 d^-1^ and 0.33 d^-1^ as a function of division rate (d^-1^). The red line indicates the growth rate range compatible with the time lag observed between BDZ and BoB (24-28 days, blue line).

## 4. Discussion

Hydrodynamics strongly regulate phytoplankton biomass, from mesoscale dispersal and dilution (Lévy et al. 2024; Lehahn et al. 2017; Ser-Giacomi et al. 2023) to the spring–neap cycles in coastal ecosystems (Cloern 1991; Blauw et al. 2012; Cadier et al. 2017) and upwelling influence (Largier 2020). Here, we focused on a single diatom species responsible for HAB. By combining field observations, transcriptomics, toxin measurements, and realistic hydrodynamic simulations, we show that physical forcing alone, primarily through dilution, and secondarily through connectivity, lead to contrasting cell physiology and bloom dynamics across neighboring coastal bays with broadly similar environmental conditions. Across the two bays, bloom development occurred within a shared seasonal framework typical of spring in the northern hemisphere marked by increasing temperature and salinity and declining nutrient concentrations. Apart from localized and transient features, abiotic conditions were similar between the two bays, yet bloom timing, magnitude, and physiological trajectories differed markedly. *P. australis* developed earlier and reached higher cell densities in the Bay of Douarnenez, while bloom progression in the Bay of Brest was delayed and more gradual. This latter pattern has been recurrently observed for many years by the French Observation and Monitoring program for Phytoplankton and Hydrology in coastal waters (REPHY) (Belin et al. 2023). The spatio-temporal offset in bloom dynamics is difficult to reconcile with spatially heterogeneous environmental forcing but emerges naturally from differences in hydrodynamic regimes.

Hydrodynamic simulations revealed dilution rates on average more than three times higher in the Bay of Brest than in the Bay of Douarnenez during the study period. Using a simple growth–dilution framework with single realistic division rates of 0.45 d^-1^ (Anderson et al. 2021) in the two bays, these differences in dilution were sufficient to reproduce the observed ∼25-day shift in bloom occurrence, implying that bloom initiation may have occurred at similar times with similar growth rates in both bays. Lower dilution in the Bay of Douarnenez would promote rapid cell density increase, while stronger dilution in the Bay of Brest would delay cell density increase. As a result, difference in dilution is a parsimonious and direct explanation to the spatio-temporal offset from the bloom onset to the bloom maximum.

The difference in dilution between the two bays may also explain why bloom termination occurs earlier in the Bay of Douarnenez. At the time of the maximum abundance in the Bay of Douarnenez, when *P. australis* strongly dominated the community, there was a sharp decrease in phosphate concentration and to a lesser extent in silicate leading to a transient increase in N:P and N:Si ratio. This probably reflected the nutrient consumption by the dense population of *P. australis* cells in a bay where both dilution and nutrient riverine inputs are low (Merceron et al., 2007; Guillam et al., 2020). This change in nutrient ratio may have triggered the bloom termination, as an increase in N:P and N:Si ratio has previously been associated with diatom bloom termination (Zhou et al. 2017; Makareviciute-Fichtner et al. 2024). In the present case, in terms of relative abundance, diatoms decreased and haptophytes and ciliates increased. In contrast, in the Bay of Brest, dilution as well as riverine inputs (Le Pape et al., 1996; Beucher et al., 2004) were stronger and cell densities increased slower, did not strongly affect nutrient concentration or ratio, and bloom termination was more progressive.

Gene expression was consistent with this hydrodynamically controlled bloom dynamic. In the two bay, a first *P. australis* gene module displayed high expression of metabolic genes, such as carbon fixation and metabolism, amino acid, fatty acid and secondary metabolites metabolism from the beginning to the maximum abundance of the bloom. This is similar to expression patterns observed in diatoms actively dividing in various other ecosystems (Marchetti et al. 2012; Gong et al. 2016; Conroy et al. 2020; Moreno et al. 2024). In the two bays, following the maximum abundance, the expression of these genes quickly decreased as cell densities dropped. A second module displayed a very strong positive correlation between gene expression and time. Interestingly, this correlation did not depend on the specific station, which is in favor of physiological transitions occurring simultaneously in the two bays. The only exception was a strong increase in expression following the maximum abundance in the Bay of Douarnenez. This module contained genes related to cell stress and mortality. In diatoms, a strong emphasis has been made on Programmed Cell Death (PCD) involving metacaspases (Bidle and Bender 2008; Graff van Creveld et al. 2021). In the present study, metacaspases were not detected in the mortality-associated expression module. Instead, it was characterized by broader functions associated with cytoskeletal dynamics, intracellular trafficking, proteostasis, energy metabolism and cellular stress. This transcriptional signature was thus indicative of severe cellular stress and potentially mortality, rather than as evidence for PCD. It would indicate that the proportion of dying cells increased regularly throughout the bloom in the Bay of Brest. In the Bay of Douarnenez, it strongly increased following maximum abundance, which may have been triggered by nutrient limitation or changes in nutrient ratios.

Domoic acid concentration per cell was often higher than 10 pg.cell-1, which is in the high range of concentration reported both in culture and in situ (usually between less than 1 to 20 pg.cell-1; Trainer et al. 2012; Ayache et al. 2019; Godinho et al. 2018; Fehling et al. 2004), although higher concentration, up to 100 pg.cell-1, has been recorded in North America (Schnetzer et al. 2007). Despite significant changes in environmental conditions linked to seasonal transition, the concentration of domoic acid per cell was not correlated with the environment. Relation between domoic acid concentration/production and available nutrients has been heavily discussed in the literature (Cochlan et al. 2023). Here we do not have the proper resolution to make a strong link between environmental conditions and DA production. In culture, an increase in extracellular domoic acid is often associated with cell lysis (Fehling et al. 2004; Ayache et al. 2019; Godinho et al. 2018). In the present study, the proportion of intracellular domoic acid was high in the bay of Douarnenez until maximum abundance was reached, then decreased rapidly. This perfectly fit with cell density as well as gene expression data suggesting that the proportion of dying cells in the population strongly increased after maximum abundance was reached. In the Bay of Brest, the proportion of intracellular domoic acid was lower, even before the maximum abundance was reached. It could indicate that domoic acid from dead or dying cells was already a significant fraction of the total domoic acid when cell densities were low and increasing in the Bay of Brest. The difference in dilution between the two bays cannot explain this pattern. Hydrodynamic connectivity provides a complementary explanation. Simulations indicate transport from the Bay of Douarnenez toward the Bay of Brest on timescales comparable to the observed bloom shift. Under this scenario, populations in the Bay of Brest would consist of a mixture of locally growing cells and older cells advected from the south. Such population heterogeneity could lead to lower intracellular DA proportion, even while total cell densities were still increasing.

Overall, our results suggest that hydrodynamic forcing, primarily through dilution, and secondarily through connectivity, acts as the dominant mechanism responsible for the spatio-temporal heterogeneity of bloom dynamics in this system. Other processes such as competition, parasitism, grazing or life cycle progression undoubtedly operate during the bloom, but our observations do not require spatial contrasts in these processes to explain the differences between bays. Difference in dilution alone is sufficient to explain a few weeks of time lag in bloom onset, one order of magnitude difference in cell densities, differences in local consumption of nutrients leading to massive shifts in nutrient ratio and profound modifications in gene expression that could explain bloom demise. In addition, directional connectivity may bring together cells with different life histories and in different physiological states. This later point needs further investigation to determine the physiological status of the transported cells, how transport affects cell physiology, and whether the transported cells may contribute or not to local growth.

Altogether, these results illustrate that hydrodynamics does not simply modulate population density and transport. It may structure the full trajectory of the bloom, from initiation to physiological decline. In numerous coastal settings, wherever bathymetry or coastline topology creates spatial contrasts in residence time or connectivity, comparable cascading effects on bloom timing, nutrient drawdown and population physiology should be expected. As a result, hydrodynamic regimes warrant as much consideration as biotic and abiotic environmental conditions to explain phytoplankton spatio-temporal dynamics.

## 6 Acknowledgments

This work is part of Lea Prigent PhD project and received financial support from the research project “MaSCoET” (Maintien du Stock de Coquillage en lien avec la problématique des Efflorescences Toxiques) financed by France Filière Pêche and the Brittany Region. The authors would like to thank the REPHY monitoring system for bloom alerts and Lauriane Baudy for help with domoic acid quantification, as well as the Ifremer RIC informatic and sebimer bio-informatic teams for support. Sequencing has been done at the GeT-PlaGe GenoToul sequencing platform (Toulouse, France).

